# New insights into antimalarial chemopreventive activity of antifolates

**DOI:** 10.1101/2021.08.17.456746

**Authors:** Chatpong Pethrak, Navaporn Posayapisit, Jutharat Pengon, Nattida Suwanakitti, Atiporn Saeung, Molnipha Shorum, Kittipat Aupalee, Kritsana Taai, Yongyuth Yuthavong, Sumalee Kamchonwongpaisan, Natapong Jupatanakul

## Abstract

Antifolates targeting dihydrofolate reductase (DHFR) are antimalarial compounds that have long been used for malaria treatment and chemoprevention (inhibition of infection from mosquitoes to humans). Despite their extensive applications, the thorough understanding of antifolate activity against hepatic malaria parasites, especially resistant parasites, have yet to be achieved. Using a transgenic *P. berghei* harboring quadruple mutant *dhfr* from *P. falciparum* (*Pb::Pfdhfr*-4M), we demonstrate that quadruple mutations on *Pfdhfr* confer complete chemoprevention resistance to pyrimethamine, the previous generation of antifolate, but not a new class of antifolate designed to overcome the resistance such as P218. Detailed investigation to pin-point stage-specific chemoprevention further demonstrated that it is unnecessary for the drug to be present throughout hepatic development. The drug is most potent against the developmental stages from early hepatic trophozoite to late hepatic trophozoite, but is not effective at inhibiting sporozoite and early hepatic stage development from sporozoite to early trophozoite. Our data shows that P218 also inhibited the late hepatic stage development, from trophozoite to mature schizonts to a lesser extent. With a single dose of 15 mg/kg, P218 prevented infection from up to 25,000 pyrimentamine-resistant sporozoites, a number equal to thousands of infectious mosquito bites. Additionally, the hepatic stage of malaria parasite is much more susceptible to antifolates than the asexual blood stage. This study provides important insights into the activity of antifolates, as a chemopreventive therapeutic which could lead to a more efficient and cost effective treatment regime.

## Introduction

Morbidity and mortality caused by malaria continue to be a heavy burden for humankind in many parts of the world. The World Malaria Report (2020) revealed more than 200 million annual cases and 400,000 annual deaths from malaria worldwide in 2019 and 2020. Antimalarial treatments have long been used to reduce the burden of malaria infection in humans; however, emergence of resistant parasites to the antimalarials currently in clinical use requires the discovery and development of new antimalarial drugs. In addition, antimalarial drugs that can only target the asexual stage are insufficient to achieve the goal of malaria elimination, because the parasite undergoes complex life cycle consisting of sexual and asexual stages in the human host and the mosquito vector. The ideal drug should be able to treat the symptoms of the disease, block the transmission from infected human to mosquito (transmission-blocking), and prevent infection from infectious mosquito bites in human (chemoprevention or prophylaxis).

The *Plasmodium* life cycle comprises multiple developmental stages in vertebrate and mosquito hosts, with each developmental stages having distinct physiology, metabolism, and function. Thus, antimalarial drugs might not be able to target more than one life stage if they do not target metabolism that are central to survival of the parasite. Malaria infection in humans occurs when *Plasmodium*-infected mosquitoes inject sporozoites into the human host during blood feeding. The injected sporozoites then infect hepatocytes in the liver where the parasites undergo endomitotic replication to generate thousands of merozoites inside hepatic schizonts over a period of one week. Once the hepatic schizonts mature, the parasites rupture to release merozoites and invade red blood cells to start erythrocytic stage, which is the stage that causes the clinical symptoms. In this stage, the parasites multiply and develop through cycles of ring, trophozoite and schizont stages to generate billions of progenies inside the human host (Cowman et al., 2016). A high replication rate is the common feature of developmental stages in different hosts, and an ideal antimalarial should target crucial factors involved in this process.

Enzymes in the folate pathway are well-defined clinically validated targets of antimalarial drugs that interfere with parasite DNA synthesis, especially the *Plasmodium* dihydrofolate reductase (DHFR). Antifolate compounds that target DHFR such as pyrimethamine (PYR) have been historically used for treatment and prophylaxis against malaria. However, the treatment has been compromised due to the mutation on DHFR of the parasite which progressively develops in *P. falciparum* with complete treatment failure of the drug in quadruple mutant (N51I+C59R+S108N+I164L) (Ahmed et al., 2006; Sirawaraporn, 1998). Recently, novel antifolate compounds have been rationally designed based on crystal structures to overcome antifolate resistance arising from DHFR mutations with P218 as a lead compound (Yuthavong et al., 2012).

Activity of antifolates as antimalarial drugs has been extensively studied only in the asexual blood stages due to the simplicity of *in vitro* drug testing assays. In contrast, antimalarial activity of antifolates against mosquito and liver stages has been investigated to a much lower extent, especially those with quadruple mutations of DHFR. This is due to the fact that *Anopheles* mosquito colonies are required for such experiments, and laboratory strains of quadruple mutant *P. falciparum* such as the V1/S strain cannot develop into gametocytes (Posayapisit et al., 2021). In the rodent malaria models, pyrimethamine resistant *P. berghei* and *P. chabaudi* strains were developed through drug pressure (Hayton et al., 2002; Nuralitha et al., 2016; van Dijk et al., 1994) but these mutations do not represent altered interactions between DHFR inhibitors and quadruple mutant *Pf*DHFR. With these limitations, there are still several knowledge gaps in the basic biology of antifolate chemopreventive activity including: 1) how much the quadruple mutations in DHFR contribute to chemoprevention resistance, 2) whether recently developed antifolates can overcome the resistance mutations in the chemoprevention setting, 3) whether it is necessary for the hepatic parasites to be exposed to the deployed antifolate throughout the hepatic development, 4) what specific hepatic developmental stages are susceptible to the antifolate, and 5) what is the antifolate level of protection against sporozoite load?

Here we used the previously developed transgenic *P. berghei* with the wild-type *Pbdhfr* replaced by quadruple mutant *Pfdhfr* (Koonyosying et al., 2020) to investigate chemoprevention activity of the previous generation as well as the new generation of antifolate compounds, represented by PYR and P218 respectively, against quadruple mutant *Pfdhfr* parasite. This study also takes an advantage of a highly potent, short half-life properties of P218 to explore stage-specific activity of antifolate compounds against hepatic parasites. This study provides important new knowledge that can further help tailor drug administration scheme to fit malaria endemicity in targeted areas.

## Materials and Methods

### Ethic statement

This study was strictly carried out in accordance to the Guide for the Care and Use of Laboratory Animals of the National Institutes of Health, and Thailand’*s* National research council. The animal protocol was approved by the BIOTEC Institutional Animal Care and Use Committee (Permit number: BT-Animal 27/2562).

### Animal and Parasite strains

Healthy 8 weeks old female ICR mice were used for all the experiments in this study. The mice were obtained from the National Laboratory Animal Center, Mahidol University, Thailand. The mice were acclimatized for at least three days before experiments and were maintained at 22±2 °C with 12/12 hour light/dark cycle. Water and food were provided *ad libitum*.

The wild-type *Pbdhfr P. berghei* ANKA strain 676m1cl1 (MRA-868) was obtained through BEI Resources, NIAID, NIH, contributed by Chris J. Janse and Andrew P. Waters. *P. berghei* with quadruple *dhfr* mutant from *P. falciparum* replacing the wild-type *dhfr* was generated in previous study (Koonyosying et al., 2020).

Laboratory strains of *Anopheles dirus* were originally collected in Mae Sod District, Tak Province, Thailand. A colony of *An. dirus* was maintained at the Department of Parasitology, Chiang Mai University and BIOTEC’s insectary at 27 °C with 70% humidity and a 12-h day/night, 30-min dusk/ dawn lighting cycle. The larvae were fed a diet of powdered fish food (Tetrabit, Germany). Adults were maintained on a 10% sucrose solution supplemented with 5% multivitamin syrup (Sevenseas, UK) *ad libitum* (Choochote & Saeung, 2013).

### *P. berghei* propagation

Asexual stage of the *P. berghei* ANKA 676m1cl1 expressing GFP-luciferase (wild-type *P. berghei dhfr*, MRA-868), and transgenic *P. berghei* ANKA *Pb*::*Pfdhfr*-4M harboring quadruple mutant *Pfdhfr* were propagated by passaging in ICR mice by intra-peritoneal (i.p.) injection of 150 μL frozen glycerol parasite stock. These parasites were passaged in mice for no longer than 8 generations without passaging in mosquito to ensure that the parasites maintain mosquito infectivity. To collect frozen parasite stocks, parasitemia in mice was monitored by thin blood smears daily until reaching 10-20%. Parasitized blood was then collected by cardiac puncture then mixed with 0.1 volume of 0.25 mM heparin and an equal volume of freezing medium (sterile 30% glycerol in PBS) then stored in liquid N_2_.

### *P. berghei* sporozoite culture

To start *P. berghei* sporozoite culture, 150 μL of frozen of *P. berghei* stock was thawed at room temperature then i.p. injected into a mouse. On the same day, a second mouse was i.p. injected with 0.2 mL of 6 mg/mL phenylhydrazine (Sigma, Cat no. P26252, USA) to stimulate reticulocyte production.

The parasitemia and gametocytemia was monitored until reaching 10-15% (from day 3-5 post infection). Infected blood was then collected using heparin as an anticoagulant and then diluted to a concentration of 5×10^7^ infected red blood cells/mL in 1X PBS buffer. The phenylhydrazine-treated mouse was then infected with 5×10^6^ infected red blood cells. Exflagellation of the parasite in the second mouse was determined by mixing 2 μL of blood collected from tail vain in 37 μL of exflagellation medium (RPMI 1640 supplemented with 25 mM HEPES (Sigma, Sigma no. H4034, USA), 2 mM glutamine (Sigma, Cat no. G7513, USA), 100 mM xanthurenic acid (Sigma, Cat. no. D120804, USA)) then incubated for 10 minutes at room temperature. Exflagellation foci were then counted under a light-contrast microscope. Mice with 10-15% parasitemia and >20 exflagellation sites per field of view of 100X magnification were used for mosquito feeding.

To prepare mosquitoes for blood feeding, female *An. dirus* were separated into waxed paper cup then starved overnight. To infect mosquitoes, infected mouse was anesthetized by 0.6 mL of 2% 2,2,2- Tribromoethanol (Sigma Cat no. T48402, USA). When the mouse was non-responsive after footpad pinching, they are placed on top of the mosquito cup, and the mosquitoes were allowed to feed for 30-45 minutes at 19±1°C. Blood-fed mosquitoes were then aspirated into a new cup then provided with 10% sucrose supplemented with 0.05% para-aminobenzoic acid (PABA, Sigma, Cat no. A9878, USA) and maintained at 19±1°C. day 8-10, GFP-fluorescent oocysts in the bloodfed mosquitoes were observed under a fluorescent microscope. Salivary glands containing sporozoites were then dissected between 20-23 days after infectious bloodmeal similar to previously published protocol (Roth, Adapa, et al., 2018; Roth, Maher, et al., 2018). Briefly, infected mosquitoes were surface-sterilized with 70% ethanol, then washed twice with 1X PBS. Salivary glands were then dissected in a drop of 1X PBS then collectively transferred to 100 μL ice-cold Schneider’s insect cell culture medium without NaHCO_3_ (Sigma, Cat no. S9895, USA). Sporozoites were then released from salivary glands by gentle grinding with sterile plastic pestle followed by pipetting for 100 times. Number of sporozoites were then counted with hemacytometer before diluting to desired number in an incomplete RPMI 1640 medium for intravenous injection.

### *In vivo* antimalarial chemoprevention assay

Chemoprevention activity of antifolates was conducted by measuring an ability of the drugs to prevent asexual stage development in mice. Briefly, antifolates such as PYR and P218 were prepared in drug carrier (0.5% w/v hydroxypropylmethylcellulose (HPMC), 0.8% v/v Tween 80 in sterile water) then orally administered following regime specific to each experiment. Sporozoites were inoculated into mice via the tail vein with the number as specified in each experiment. Drug carrier (0.5% HPMC and 0.4% tween-80 in water) was used as an untreated control of each experiment. To determine asexual stage development, Giemsa-stained thin blood smear of each mouse was monitored between 4 to 14 after sporozoite injection.

## Results

### Quadruple mutation on *P. falciparum dhfr* gene renders PYR ineffective for chemoprevention

We utilized a transgenic *P. berghei* harboring *P. falciparum dhfr*-4M developed by our group (Koonyosying et al., 2020) to determine chemoprevention activity of PYR against quadruple mutant parasite. We have confirmed that the transgenic *Pb*::*Pfdhfr*-4M parasite still maintains an ability to develop into mosquito and liver stages (Figure 1).

**Figure 1.**
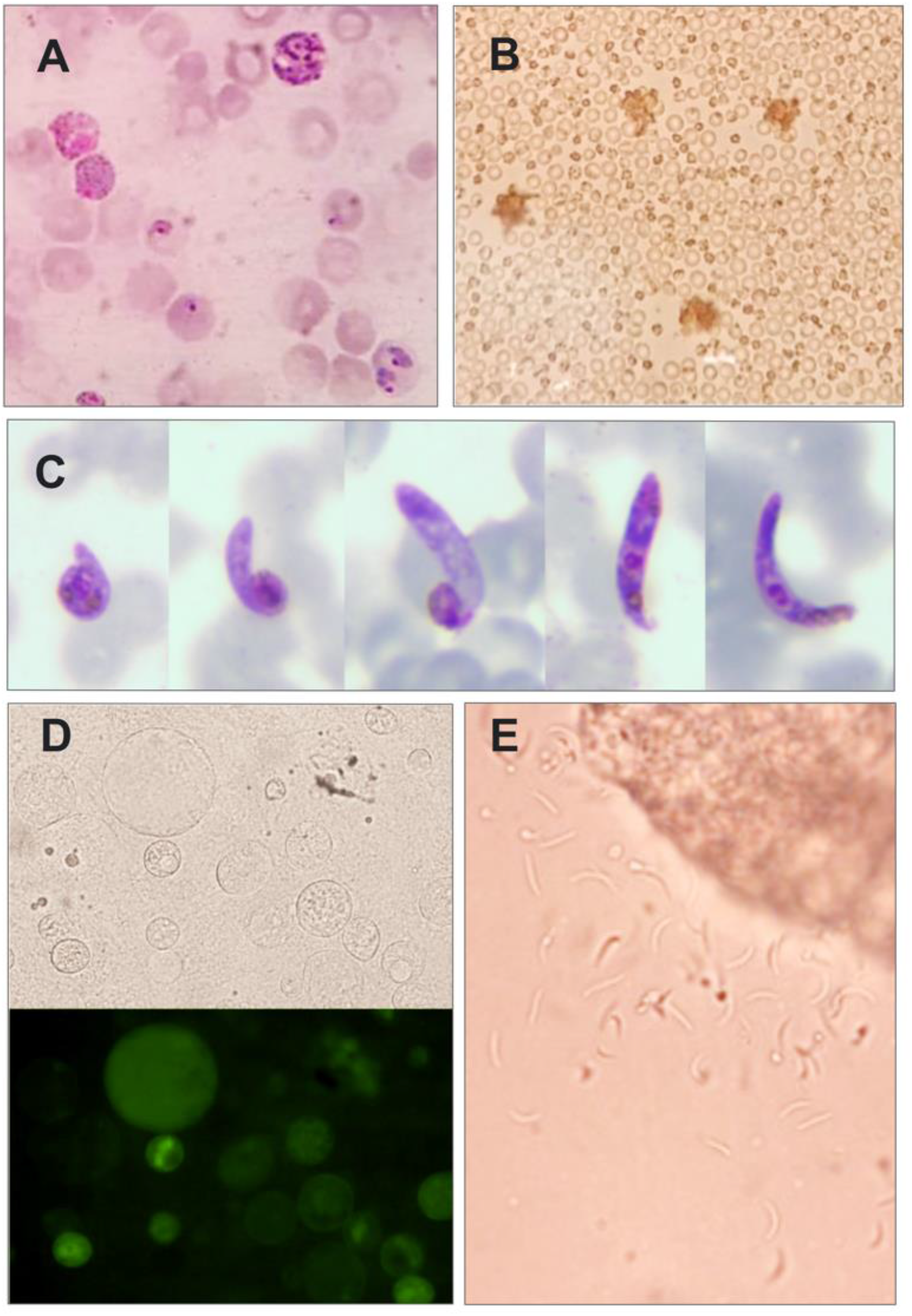
The transgenic parasite maintains an ability to develop to mosquito stages. A) Mature male and female gametocytes, B) male gametocyte exflagellation under bright field microscope, C) Giemsa-stained ookinete, D) Midgut oocysts under bright field and fluorescent microscope, and E) sporozoites released from squashed salivary gland under bright-field microscope.

First, two doses of 30 mg/kg PYR were administered one day before and one day after intravenous injection of 2,000 wild-type or transgenic *Pb*::*Pfdhfr*-4M sporozoites. Using this administration scheme, the sporozoites were exposed to the drug as soon as they entered the circulatory system, and also after hepatocyte invasion. All mice in the control (carrier) group became infected by either parasite following the challenge (Figure 2). While 30 mg/kg PYR could inhibit wild-type *P. berghei*, the drug failed to prevent infection by transgenic *Pb*::*Pfdhfr*-4M (Figure 2). These results demonstrated that the quadruple mutations in *Pfdhfr* gene causes complete chemoprevention resistance to PYR, the first generation of antifolate.

**Figure 2.**
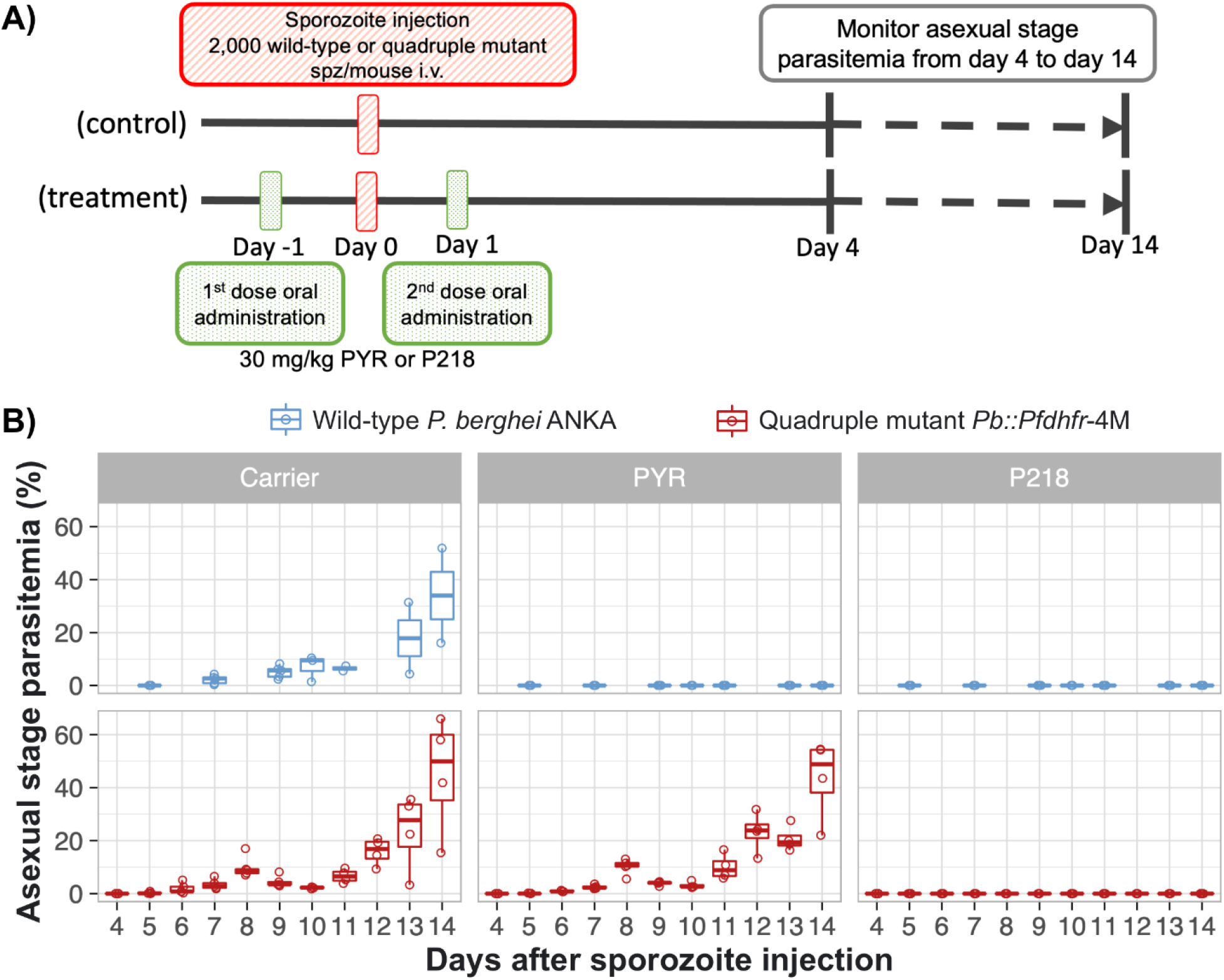
Quadruple mutation on *Plasmodium falciparum dhfr* gene confers complete resistance to previous generation antifolate. A) Schematic diagram of *in vivo* chemoprevention assay to test activity of antifolates against wild-type and quadruple mutant *Pfdhfr* transgenic parasite. Mice in the treatment group were orally treated with two doses of 30 mg/kg PYR or P218 one day before and one day after sporozoite injection. Mice in the control group were given with carrier for drug administration using the same scheme as the treatment group (0.5% w/v hydroxypropylmethylcellulose, 0.8% v/v Tween 80 in sterile water). B) Asexual stage parasitemia of the wild-type *P. berghei* (ANKA) and the transgenic *Pb::Pfdhfr*-4M during 4-14 days after sporozoite challenge (intravenous injection with 2,000 sporozoites).

### Novel antifolate potently inhibits liver stage of quadruple mutant parasite

Using P218 as a model for a new class of antifolates, subsequent experiments established that this lead compound can inhibit liver stage infection of both the wild-type and quadruple mutant parasites. Oral administration of 30 mg/kg P218 on day −1 and day 1 relative to sporozoite challenge prevented asexual stage infection in both the wild-type and the transgenic PYR-resistant parasites (Figure 2).

After chemoprevention activity of P218 against the wild-type and quadruple mutant parasite was confirmed at 30 mg/kg, we then further investigate how much quadruple mutations alter parasite sensitivity to P218 using various doses of the drug.

In the wild-type parasite, P218 could completely prevent parasite development into blood stage when orally treated at 0.5 mg/kg (Figure 3). Although complete blocking of infection was not achieved, the lower doses of P218 could improve survival rate at 14 dpi. At 0.125 mg/kg, none of the mice died at 14 dpi, and 2 out of 4 mice became infected (Figure 3). These results demonstrate that although P218 at the dose lower than 0.5 mg/kg was not able to completely prevent parasite development into blood stage, the drug could still reduce the burden of malaria.

**Figure 3:**
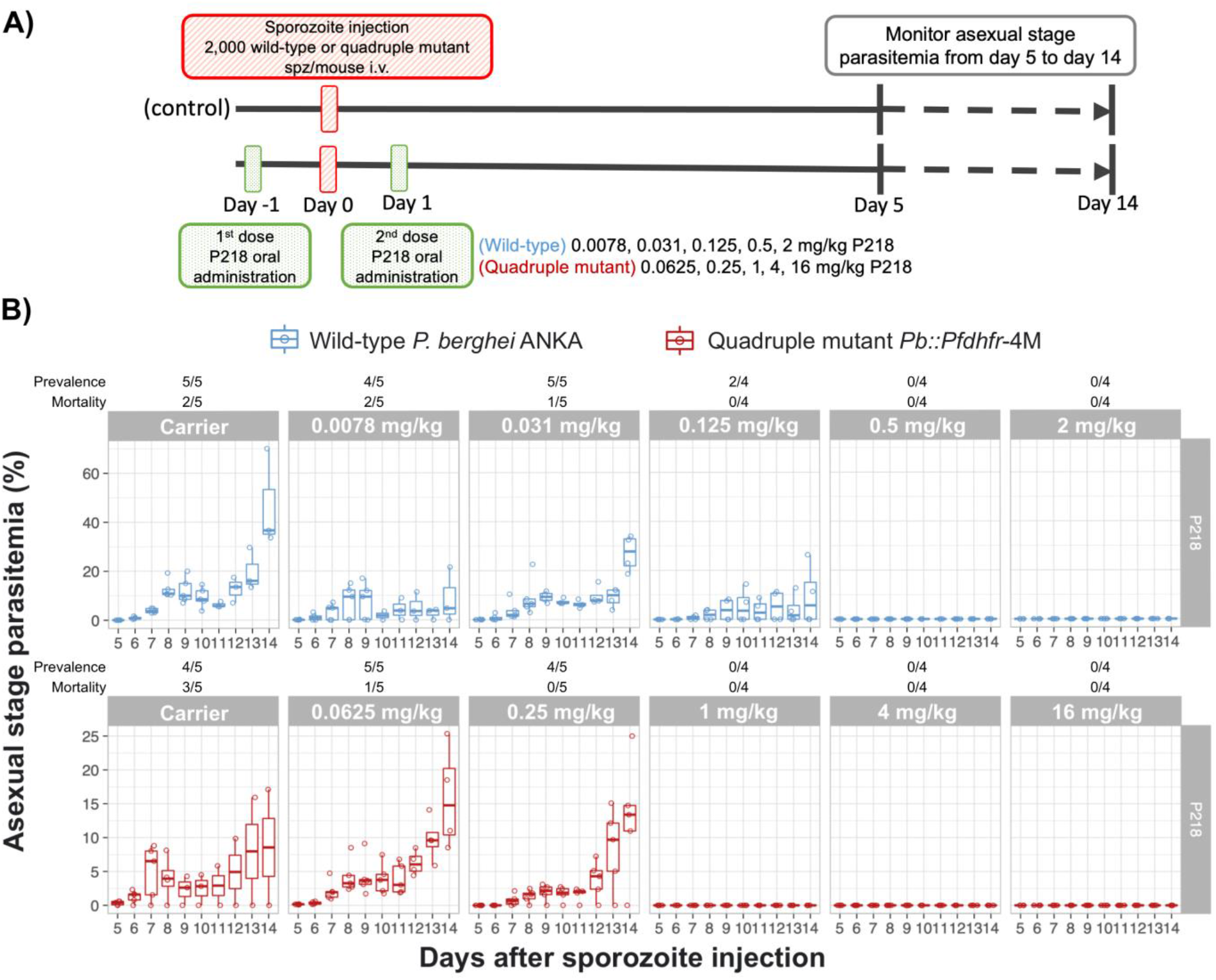
Rationally designed antifolate, P218, is highly potent for chemoprevention against both wild-type and PYR-resistant parasites. A) Schematic diagram of *in vivo* dose-response analysis for chemoprevention against wild-type and quadruple *Pfdhfr* mutant transgenic parasites. Mice infected by the wild-type *P. berghei* ANKA were orally treated with 2 doses of 0.0078, 0.031. 0.125, 0.5, or 2 mg/kg P218 while those infected with PYR-resistant parasites were orally treated with 2 doses of 0.0625, 0.25. 1, 4, or 16 mg/kg P218. The control groups were treated with carrier used for oral drug delivery (0.5% w/v hydroxypropylmethylcellulose, 0.8% v/v Tween 80 in sterile water) B) Asexual stage parasitemia of the wild-type *P. berghei* (ANKA) and the transgenic *Pb::Pfdhfr*-4M during 5-14 days after sporozoite challenge and treatment with various doses of P218.

P218 is potent for chemoprevention against the PYR-resistant parasites. At an oral dose of 1 mg/kg, P218 could completely prevent transgenic PYR-resistant *P. berghei* development into blood stage in all mice (Figure 3). The lower doses at 0.25 and 0.0625 mg/kg reduced the mortality at 14 dpi demonstrating that these lower doses could reduce burden from PYR-resistant malaria although complete blocking was not achieved. This minimum inhibitory concentration of P218 against PYR-resistant parasite is only 2-fold higher than those of the PYR-sensitive parasite suggesting that the rationally designed antifolate can overcome resistance from quadruple mutation in *Pfdhfr* and is thus effective for chemoprevention against the prevalent resistant parasites in the field.

### Development from hepatic ring to trophozoite is the stage the most susceptible to antifolate

Characteristics of P218 including high potency, rapid absorption (Tmax in mice = 0.25 hour), and short half-life (oral half-life in mice = 4 hours) (Yuthavong et al., 2012), allow us to dissect specific hepatic developmental stage that is the most susceptible to antifolate. When administered at minimum inhibitory dose, the compound should enter the circulation almost immediately and quickly cleared from the body thus exposing only to that specific developmental stage. Previous studies showed that *P. berghei* develops from sporozoite to ring stage during the first day, ring to late trophozoite stage during the second day, and late trophozoite to schizont during the third day (Figure 4) (Ng et al., 2015). In the following experiment, a single dose of P218 was administered at the minimum inhibitory concentration (1 mg/kg) on various days from a day before sporozoite injection to three days after infection.

**Figure 4:**
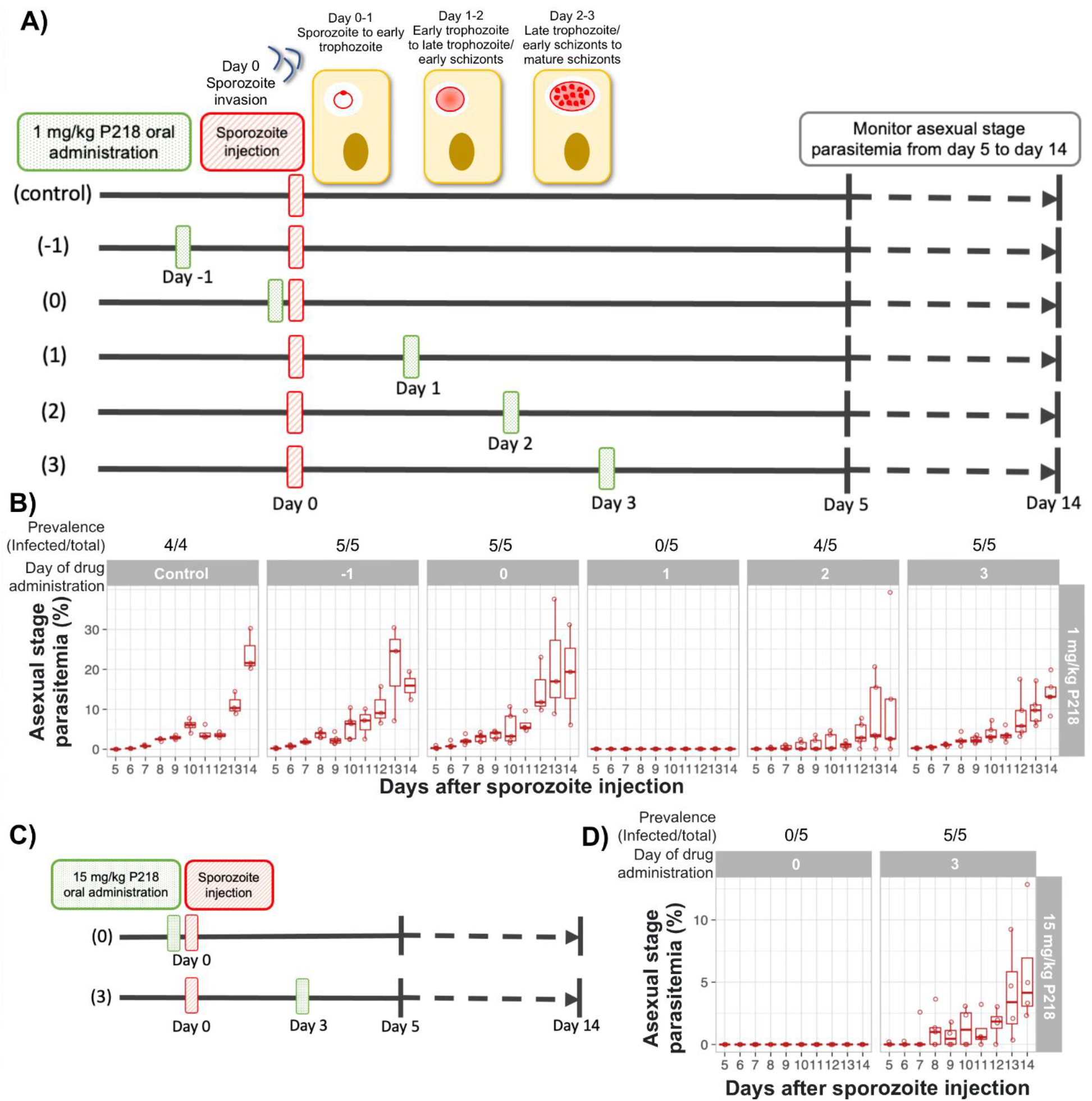
Development from early trophozoite to early schizonts is the hepatic stage most susceptible to antifolate. A) Schematic diagram of an *in vivo* experiment to determine hepatic stage most susceptible to antifolate. Mice were orally treated with a single dose of 1 mg/kg P218 on different days from day −1 to day 3 relative to sporozoite challenge. The oral administration on day 0 was conducted three hours before sporozoite challenge. All mice were intravenously infected with 2,000 transgenic *Pb::Pfdhfr*-4M sporozoites. B) Asexual stage parasitemia of the transgenic *Pb::Pfdhfr-4M* during 5-14 days after sporozoite challenge and treatment with a single dose of 1 mg/kg P218 on day −1 to day 3 relative to sporozoite challenge. C) Schematic diagram of an *in vivo* experiment to determine the effect of an increased dose (15 mg/kg) on early and late hepatic stage parasite. D) Asexual stage parasitemia of the transgenic *Pb::Pfdhfr-4M* during 5-14 days after sporozoite challenge and treatment with a single dose of 15 mg/kg P218 on day 0 (3 hours before) or day 3 relative to sporozoite challenge.

**Figure 5:**
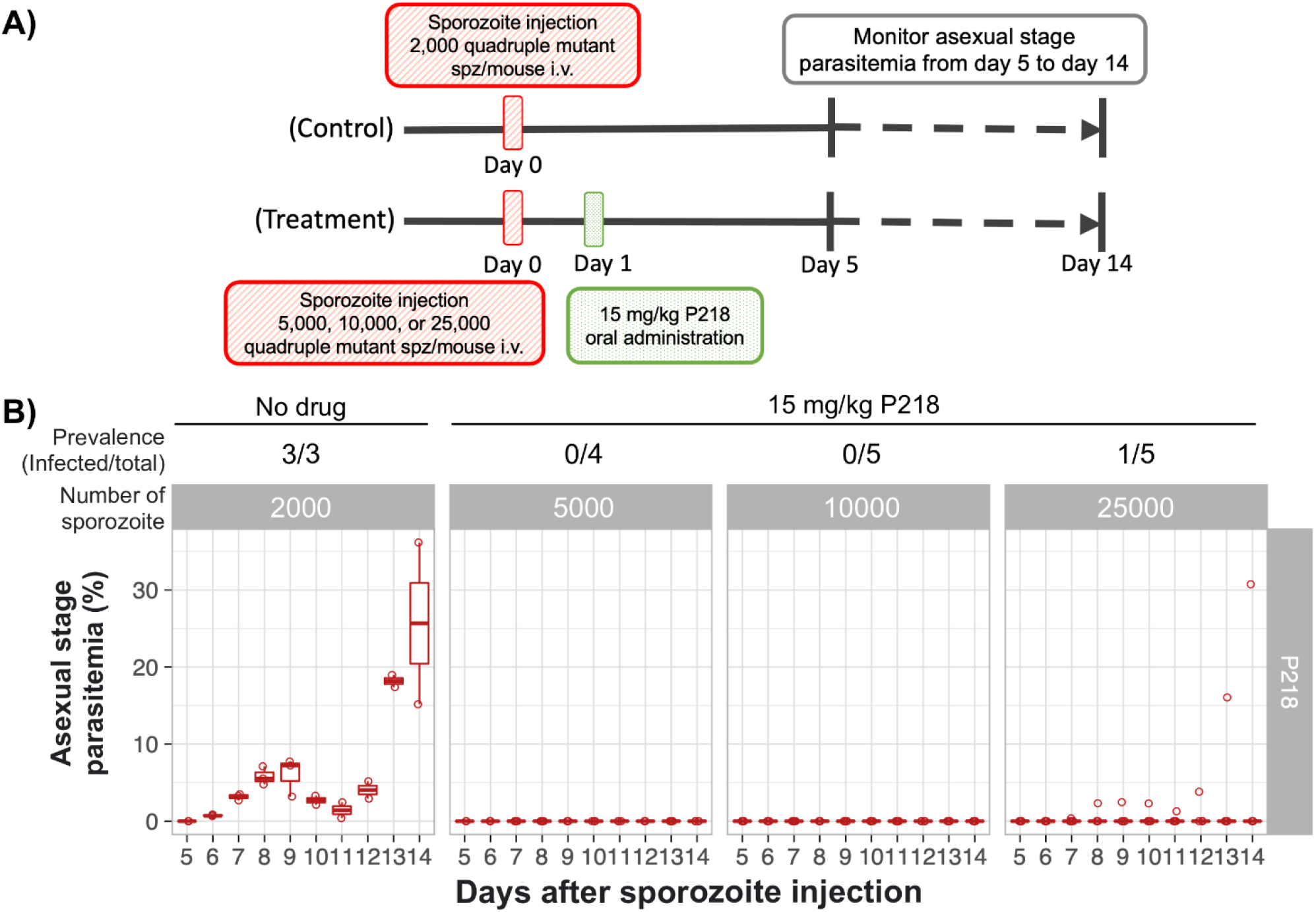
P218 prevents infection by a large number of PYR-resistant sporozoites. A) Schematic diagram of an *in vivo* experiment to determine the number of PYR-resistant sporozoite that 15 mg/kg P218 can completely prevent infection. Mice were intravenously infected with 5,000, 10,000 and 25,000 sporozoites then orally treated with 15 mg/kg P218 1 day after the challenge. The control group was infected with 2,000 sporozoites then carrier used for oral delivery was administered 1 day after the challenge. B) Asexual stage parasitemia of the transgenic *Pb::Pfdhfr-4M* during 5-14 days after a challenge with 5,000, 10,000, or 25,000 sporozoites followed by an oral dose of 15 mg/kg on the following day.

A single oral dose of 1 mg/kg P218 was able to completely clear the *P. berghei* liver stage infection, with no recrudescence of parasites (monitored with thin blood smear until day 28 post-sporozoite injection) when administered on day 1 (Figure 4), demonstrating that a single dose of P218 is sufficient for chemoprevention when administered with this scheme. The treatment with single dose of P218 on day 1 resulted in a complete prevention of parasite infection similar to the treatment with two doses in Figure 3.

Administration of the compound one day before (day −1) or three hours before (day 0) had no impact on liver stage infection both in term of infection prevalence and intensity (Figure 4). Although all mice became infected when P218 was administered two or three days after sporozoite challenge, lower parasitemia and slower parasite development into erythrocytic stage were observed (Figure 4). These results suggested that the hepatic developmental stage during one day after sporozoite challenge, namely from ring to late trophozoite, is the most susceptible to P218.

The maximum dose of P218 used during clinical development in human was 1,000 mg, which could be translated into 13-14 mg/kg (Chughlay et al., 2020, 2021). Therefore, we next investigated if a single oral administration of 15 mg/kg P218 could improve the chemoprevention when the compound was treated on the non-optimal days (same day and three days after sporozoite challenge). Although the increased dose of P218 administered three days after sporozoite challenge could not improve chemoprevention compared with the treatment at 1 mg/kg, the increased dose prevented infection of all the mice when administered on the same day of sporozoite challenge (Figure 4). These results demonstrates that the increased dose of P218 could prolong the chemoprevention activity of P218 administered earlier than the optimum time and emphasize that chemoprevention activity of antifolates is the most active against the intrahepatic ring to trophozoite developmental stage.

### P218 could prevent infection by a large number of PYR-resistant sporozoites

Information on how many sporozoites the chemopreventive drug can protect against, will be beneficial to tailor drug administration schemes in malaria endemic regions and determine the protection in relation to the number of potential infectious bite of the endemic areas. In the subsequent experiment, mice were infected with 5,000; 10,000; and 25,000 transgenic PYR-resistant *P. berghei* sporozoites to determine if P218 is potent enough in chemoprevention against high number of PYR-resistant sporozoites. With the single oral dose of 15 mg/kg, P218 completely prevent infection in all mice infected up to 10,000 transgenic sporozoites. In the group injected with 25,000 transgenic sporozoites, no blood stage parasites were detected in four out of five mice suggesting a borderline limit of chemoprevention at this number.

## Discussion

The resistance to antifolates such as pyrimethamine has long been a problem that started soon after the drug was introduced many decades ago. A recent study demonstrated that the resistant parasites persist in the malaria endemic area even after the long-term withdrawal of antifolate use (Sugaram et al., 2020). Although the hepatic and blood stage development follow the same pattern from rings to trophozoites and schizonts, the hepatic schizogeny contains many more rounds of DNA replication and resulted in a much larger number of merozoites (Matthews et al., 2018). It is possible that this might lead to a difference in susceptibility to antifolate between hepatic and blood stages, especially in the antifolate resistant parasites.

This study sought to characterize antifolates as chemopreventive antimalarial drugs, especially against the quadruple mutant *Pfdhfr* parasite. Using the transgenic *P. berghei* harboring quadruple mutant *Pfdhfr* in place of the wild-type *Pbdhfr* as an *in vivo* antifolate resistant model, we demonstrated that the quadruple mutation on *Pfdhfr* does not only leads to PYR treatment failure (Koonyosying et al., 2020) but also confers complete resistance to PYR chemoprevention. Antifolates of the previous generation are then no longer effective for both applications in most malaria endemic areas, thus prompting the development of alternative treatments. The benefit of having a well-known target allows characterization of resistance mechanism and rational design of new classes of antifolates that can overcome the resistance (Yuthavong et al., 2012). Although the new class of antifolates such as P218 can inhibit asexual stage and male gametocyte exflagellation of quadruple mutant parasites (Koonyosying et al., 2020; Posayapisit et al., 2021; Yuthavong et al., 2012), the information on how potent these new compounds are for the chemoprevention activity against quadruple mutant parasite has never been experimentally demonstrated due to the lack of reliable liver stage model of the resistant parasite.

Our current study demonstrates a high potency of P218 against the PYR-resistant parasite both in terms of drug concentration required to completely inhibit the liver stage parasite as well as the number of sporozoite it can inhibit. In our model, the quadruple mutation of *Pfdhfr* gene resulted in an approximately two-fold increase in P218 minimum inhibitory dose required for complete chemoprevention from 0.5 mg/kg to 1 mg/kg suggesting that the rationally designed compound can overcome resistance in chemoprevention. It should be noted that our experiments used the wild-type parasite carrying *P. berghei dhfr* not the wild-type *P. falciparum dhfr*. However, previous studies showed that P218 has comparable blood stage inhibitory concentration against the wild-type *P. berghei* (0.9 nM, (Swann et al., 2016)) and wild-type *P. falciparum* (1 nM, Medicines for Malaria Venture, personal communications). Therefore, we speculate that the chemoprevention minimum inhibitory dose obtained from the wild-type *P. berghei* can be a representative of parasite harboring wild-type *Pfdhfr*.

Detailed understanding of the chemoprevention activity of antifolate allows optimization of the drug administration scheme for the most cost effective and efficient malaria control for malaria endemic areas. P218 administration at three hours before sporozoite injection exposes two parasite stages to this potent antifolate compound: sporozoites which were immediately exposed to this potent antifolate compound in the circulation, and early hepatic parasite during the development from sporozoite to hepatic ring stage. Despite these exposures during the first day, the parasite could complete intra-hepatic development and the outcome of infection was not different from the untreated group suggesting that antifolate compounds are not active against sporozoite and very early intra-hepatic stages. During approximately the first 20 h after the invasion of the hepatocyte, *P. berghei* develops into hepatic ring and early-trophozoite, the parasites remain haploid with single nucleus in G1 phase (Graewe et al., 2012). Since antifolate compounds target the pathway involved in DNA synthesis, these compounds cannot inhibit these non-replicative early hepatic stages.

Around 20 hours after sporozoite invasion into hepatocytes, the parasites develop into trophozoites as the cell cycle stage goes from G1 to M1 phase. During trophozoite development, each single parasite continue DNA synthesis and nuclear division to generate 20,000 to 30,000 new parasites in approximately 30 hours (Graewe et al., 2012). Consequently, DNA synthesis and nucleotide metabolism is crucial for this developmental stage. P218 treatment at one day (24 hours) after *P. berghei* sporozoite injection completely inhibited parasite development into blood stage, confirming that the development from trophozoite to schizonts (schizogeny) stage is extremely susceptible to antifolate due to their highly replicative nature. Interestingly, although P218 was cleared from the circulation in the following days due to its short half-life, no blood stage parasite was detected in all the treated mice. This suggests that if the parasite cannot accumulate enough DNA, they cannot stay dormant and quickly perish. This might be due to the parasite undergoing several check points prior to schizogeny, including S and G2 checkpoints (check for successful DNA replication or DNA damage) as well as M checkpoint (check for chromosome attachment to the spindle) (Matthews et al., 2018).

Once the hepatic schizogeny is completed or almost completed, the antifolate no longer affects the parasite’s ability to further progress to blood stages as demonstrated by the fact that P218 could no longer inhibit the matured hepatic parasite even at an increased dose (15 mg/kg). Due to a much higher number of parasites to be inhibited after progression to blood stage, multiple doses of P218 is needed for malaria treatment as demonstrated in previous study (Chughlay et al., 2021; Koonyosying et al., 2020).

It is interesting to note that folate pathway is also involved in other biological process such as methionine synthesis, but the compounds could not inhibit specific developmental stage with high protein synthesis such as the development from ring to trophozoite. Any other antifolate inhibitory activity against other biological processes during hepatic development, is likely to be negligible to be considered as additional mode of action of the compounds. When compared to previous stage-specific activity of antifolates against blood stage, our study coincides with previous findings (Dieckmann & Jung, 1986) that antifolates mainly target the parasites’ DNA synthesis during blood stage development.

This study provides insights into the nature of chemoprevention of P218 that compliment data from a P218 clinical trial on sporozoite challenge (Chughlay et al., 2021) that maybe beneficial in improving drug regime in clinical use. First, the Chughlay et al. clinical trial (2021) stated that it was unclear how much chemoprevention activity of P218 was due to activity against liver stage vs activity against the blood stage parasites. Our stage-specific experiment clearly demonstrated that the activity against the hepatic parasite is the main contributor to the chemoprevention. Second, our study demonstrated that P218 is highly active against the quadruple mutant parasite thus providing a data to support its potential use in the real-world where resistant parasites are predominant. Third, this study confirms that antifolates have superior potency against liver stage compared to the blood stage. We previously demonstrated that 10 mg/kg P218 was required to completely clear asexual stage parasite using the standard 4-day suppressive test (Koonyosying et al., 2020) while only a single dose of 1 mg/kg P218 is required for complete chemoprevention. It is possible that the higher sensitivity is due to a combination of more rounds of endomitosis during hepatic development and the lower number of parasites to be inhibited. Fourth, P218 plasma concentration required for chemoprevention in the real-world setting might be overestimated due to the number of sporozoites used to challenge in most preclinical and clinical studies. Our study demonstrated a correlation between dose required for chemoprevention and the number of sporozoites during the infection, which 1 mg/kg P218 could prevent infection from 2,000 sporozoites and 15 mg/kg P218 could prevent infection from 25,000 sporozoites. During transmission from infected mosquitoes to humans, the majority of mosquitoes ejected on average ten sporozoites per bite (Graumans et al., 2020). Thus, it is likely to require even lower doses for complete chemoprevention.

Lastly, the stage-specific experiments agrees with the clinical trial that the longer exposure of P218 throughout hepatic development is not necessary to maintain chemoprevention. In the real-world setting, it is impossible to predict when each person will be bitten by infectious mosquitoes. When translated into clinical use, the candidate drug might be administered at longer intervals that have potential to target the development from hepatic ring to early schizonts. In addition, since P218 is highly effective against the parasite and does not require high plasma concentration for chemoprevention, it is reasonable to administer P218 on every two to three days based on its half-life of 11-20 hours in human (Chughlay et al., 2020, 2021). In addition, it will be of greater benefit to develop a slow-release formulation that can sustain moderate levels of the candidate drug overtime.

One interesting additional application of new antifolates, especially P218, is to use these drugs to assist immunization by attenuated sporozoites, which has proved to provide a long-lasting immunity against malaria thus offering a potential vaccine candidate. The attenuation of sporozoites have been achieved by irradiation or genetic modification but previous reports also demonstrated a potential to use chemopreventative drugs such as primaquine or pyrimethamine to attenuate the hepatic parasite (Friesen et al., 2011). Since the P218 is safe and highly potent in suppressing a large number of liver-stage parasites, it could potentially be used in such application. Our results demonstrated that P218 did not kill sporozoites but rather inhibit development of hepatic stages. Consequently, parasites are still able to invade hepatocytes and attenuated during intrahepatic development. These attenuated parasites in infected cells are likely to be processed through the antigen presentation by class-I major histocompatibility complex (MHC) and triggers cytotoxic T-cell responses, an important part in antimalarial immunity (Kurup et al., 2019). It will be interesting to compare immune responses between irradiated sporozoites as well as other spotozoite-targeted vaccines to the infection by active sporozoites that was rescued by P218 treatment.

## Acknowledgements

This work was financially supported by Research Chair Grant (P-1850116) from the National Science and Technology Development Agency (NSTDA), Thailand to SK, and BIOTEC Research unit director initiative grant to NJ (P-1851424). P218 compound was kindly provided by Medicine for Malaria Venture (MMV). *P. berghei* ANKA 676m1cl1 expressing GFP-luciferase (MRA-868), contributed by Chris J. Janse and Andrew P. Waters was obtained through BEI Resources, NIAID, NIH. We also thank Dr. Warangkhana Songsungthong for assistant with the initial set up of insectary and *P. berghei* culture. We are grateful to Asst. Prof. Dr. Patchara Sriwichai from Department of Medical Entomology, Faculty of Tropical Medicine, Mahidol University, who kindly provided *An. dirus* adult used to establish the colonies used in this study to A. Saeung. We also thank Dr. Jose ? L. Ramirez from USDA for editorial assistance with the manuscript.

